# Whole-genome sequence analysis reveals evolution of antimicrobial resistance in a Ugandan colistin resistant *Acinetobacter baumannii*

**DOI:** 10.1101/2020.06.18.159236

**Authors:** Dickson Aruhomukama, Ivan Sserwadda, Gerald Mboowa

**Author notes:** Correspondence: Gerald Mboowa.

## Abstract

In recent times, pan-drug resistant *Acinetobacter baumannii* have emerged and continue to spread among critically ill patients, this poses an urgent risk to global and local human health. This study sought to provide the first genomic analysis of a pan-drug resistant *Acinetobacter baumannii* from Uganda and Africa, and to tell a story of mobile genetic element-mediated antibiotic resistance evolution in the isolate. It was an in-silico study in which intrinsic and acquired antibiotic resistance genes, and/or chromosomal resistance mutations were identified using PATRIC, CARD, NDARO and ResFinder. Screening for insertion sequences was done using ISfinder. Also, plasmid screening, phylogenetic analysis and sequence typing were performed using PlasmidFinder, PATRIC and Gubbin, and MLST respectively.

The isolate belonged to the Sequence type 136, belonging to Clonal complex 208 and Global complex 2. This isolate shared close homology with strains from Tanzania. Resistance in the isolate was chromosomally and mobile genetic element-mediated by *Acinetobacter*-derived cephalosporinases and carbapenem hydrolyzing class D β-lactamses, *bla*_*OXA-2, 51, 5 88, 317*_, *bla*_*ADC-2, 25*_. Colistin resistance was associated with previously documented mutants, *lpxA* and *lpxC*. Other key resistance genes identified were: *aph(3”)-lb, aph(6)-ld, aph(3’)-la, aac(3)-lld, aac(3)-lla, aph(3’)-l, aph(3”)-l, aph(6)-lc, aph(6)-ld, aac(3)-II, III, IV, VI, VIII, IX, X, macA, macB, tetA, tetB, tetR, dfrA*, and those of the *floR* family. RSF1010 like IncQ broad-host-range plasmids and features of pACICU1, pACICU2, and p3ABAYE *Acinetobacter baumannii* plasmids namely partitioning proteins ParA and B were present. Insertion sequences present included IS*3*, IS*5*, IS*66* and those of the ISLre*2* families.

The study described for the first time a pan-drug resistant *Acinetobacter baumannii* from Uganda, and told a story of mobile genetic element-mediated antibiotic resistance evolution in the isolate despite being limited by pan-drug resistance phenotypic data. It provides a basis to track trends in antibiotic resistance and identification of emerging resistance patterns in *Acinetobacter baumannii* in Uganda.

## Background

Current times have witnessed the emergence and continuous spread of pan-drug resistant *Acinetobacter baumannii* among critically ill patients, this poses an urgent risk to global human health^1^. The World Health Organisation (WHO) in its 2017 report on “priority pathogens – a catalog of 12 families of bacteria that pose the greatest threat to human health” categorized *Acinetobacter baumannii* in priority 1 or the critical category^1^. *Acinetobacter baumannii* is a Gram-negative opportunist pathogen with a high incidence in immunocompromised individuals, particularly those attending intensive-care units^2^. This organism exhibits an extraordinary ability to upregulate or acquire antibiotic resistance genes, a characteristic that enhances its spread and, makes it one of the bacteria that threatens the current antibiotic era^2^. However, limited information remains on the genetic antimicrobial resistance determinants as well as the mobile genetic element-mediated antibiotic resistance evolution in *Acinetobacter baumannii* especially from Uganda. This work sought to fill this gap by providing the first genomic analysis of a pan-drug resistant *Acinetobacter baumannii* from Uganda, and to tell a story of mobile genetic element-mediated antibiotic resistance evolution in the isolate.

## Materials and methods

### Isolate metadata and source of whole-genome sequences

*Acinetobacter baumannii* SRR8291681_Uganda whole-genome sequences were obtained from the NCBI’s Sequence Read Archive (SRA) repository, these had been submitted in December 2018 by J. Craig Venter Institute (JCVI) under a study entitled the whole-genome sequencing of carbapenem-resistant Gram-negative clinical isolates from Mbarara Regional Referral Hospital, Western-Uganda and BioProject: PRJNA508495 (https://www.ncbi.nlm.nih.gov/sra/SRR8291681/). In addition, 8 other available *Acinetobacter baumannii* whole-genome sequences for comparison were purposively obtained from the same source, and these were from; Tanzania, South Korea, United States, Canada, Germany, and Sweden. Mbarara Regional Referral Hospital is located in Mbarara Municipality, which is 286 km south west of Kampala the capital city of Uganda. It is a public hospital funded by the Government of Uganda through Ministry of Health. It is the referral hospital for south western Uganda serving 10 districts with a population of more than 2.5 million people. It also receives patients from neighbouring countries of Rwanda, Tanzania and Democratic Republic of Congo.

### Whole-genome sequencing

The whole genome of *Acinetobacter baumannii* SRR8291681_Uganda was sequenced using the Illumina NextSeq 500.

### Read quality assessment, genome assembly and annotation

FastQC v0.11.8 (https://www.bioinformatics.babraham.ac.uk/projects/download.html) was used to assess the quality of the reads. *De novo* genome assembly of the Illumina NextSeq 500 reads from *Acinetobacter baumannii* SRR8291681_Uganda was performed using the Unicycler v0.4.8-beta^52^ with SPAdes v3.11.1^53^, with default parameters. Following genome assembly, gene calling and automatic functional annotation were done for all the contigs using the Pathosystems Resource Integration Center (PATRIC 3.5.36) identifying 4,037 coding regions on the chromosome with 64 tRNA and 3 rRNA. PATRIC uses the RAST tool kit to provide an annotation of genomic features in default parameters (https://www.patricbrc.org/).

### Identification of antibiotic resistance genes

The identification of antibiotic resistance genes from the complete Illumina NextSeq 500 assembly was performed using PATRIC 3.5.36. Additional screening for antibiotic resistance genes was performed by comparison (BLASTp; sequence identity>=40%; E-value<=0.0001) of all predicted coding regions against the PATRIC’s Antibiotic Resistance Genes Database, the Comprehensive Antimicrobial Resistance Database (CARD)^54^, and the National Database of Antibiotic Resistant Organisms (NDARO) (https://www.ncbi.nlm.nih.gov/pathogens/antimicrobial-resistance/). Furthermore, acquired antibiotic resistance genes and/or chromosomal mutations were identified using ResFinder version 3.1 (https://cge.cbs.dtu.dk/services/ResFinder/) at a threshold for percentage identity of 90% and a minimum length of 60%. Manual inspection of the antibiotic resistance genes was performed to improve their functional annotation based on known and correct start sites as well as to identify point mutations which may contribute to a resistant phenotype. Resistance gene loci were screened for known insertion sequences by comparison against the ISFinder database set at a minimum % coverage of 70% and a threshold for minimum % identity of 30% (https://isfinder.biotoul.fr/). Finally, screening for the presence of plasmids using PlasmidFinder 2.0 set at 60% minimum % coverage and the threshold for minimum % identity 95% (https://cge.cbs.dtu.dk/services/PlasmidFinder/) was done.

### Phylogenetic analysis and multilocus sequence typing (MLST)

The building of the phylogenetic tree was performed using the PATRIC server with only 1,000 genes selected randomly from the assembled contigs and default parameters including no genomic duplications and insertions. Multilocus sequence typing was performed using MLST v2.0 (https://cge.cbs.dtu.dk/services/MLST/). The results of phylogenetic analysis obtained from the the PATRIC server were verified using Genealogies Unbiased By recomBinations In Nucleotide Sequences (Gubbins) (https://sanger-pathogens.github.io/gubbins/), this iteratively identifies loci containing elevated densities of base substitutions while concurrently constructing a phylogeny based on the putative point mutations outside of these regions enabling more accurate, recombination free phylogenetic analysis of *Acinetobacter baumannii* genomes known to be full of regions acquired as a result of recombination events.

## Results

### Read quality assessment

FastQC v0.11.8 was used to assess the quality of the reads, *Acinetobacter baumannii* SRR8291681_Uganda was found to have a coverage of >30X, generated 6.19 million paired-end 150 bp genomic reads. The assembled genome of *Acinetobacter baumannii* SRR8291681_Uganda has N50 and L50 of 337442 and 5 respectively.

### General genome features of *Acinetobacter baumannii* SRR8291681_Uganda

SRR8291681_Uganda consists of a circular chromosome spanning 4173379 base pairs in length with an average G+C content of 38.95%. The genome quality was generally good, with coarse and fine consistencies accounting for 98.7% and 97.6% respectively. The chromosome of the isolate shares close homology with *Acinetobacter baumannii* ERR1989115 and ERR1989084 strains originating from Tanzania (**Figure 1**). Multilocus sequence typing (MLST) revealed that SRR8291681_Uganda belonged to the previously reported sequence type (ST) 136^3^, this ST have been documented to belong to clonal complex 208 (CC208) and global complex 2 (GC2)^4^.

**Figure 1.**
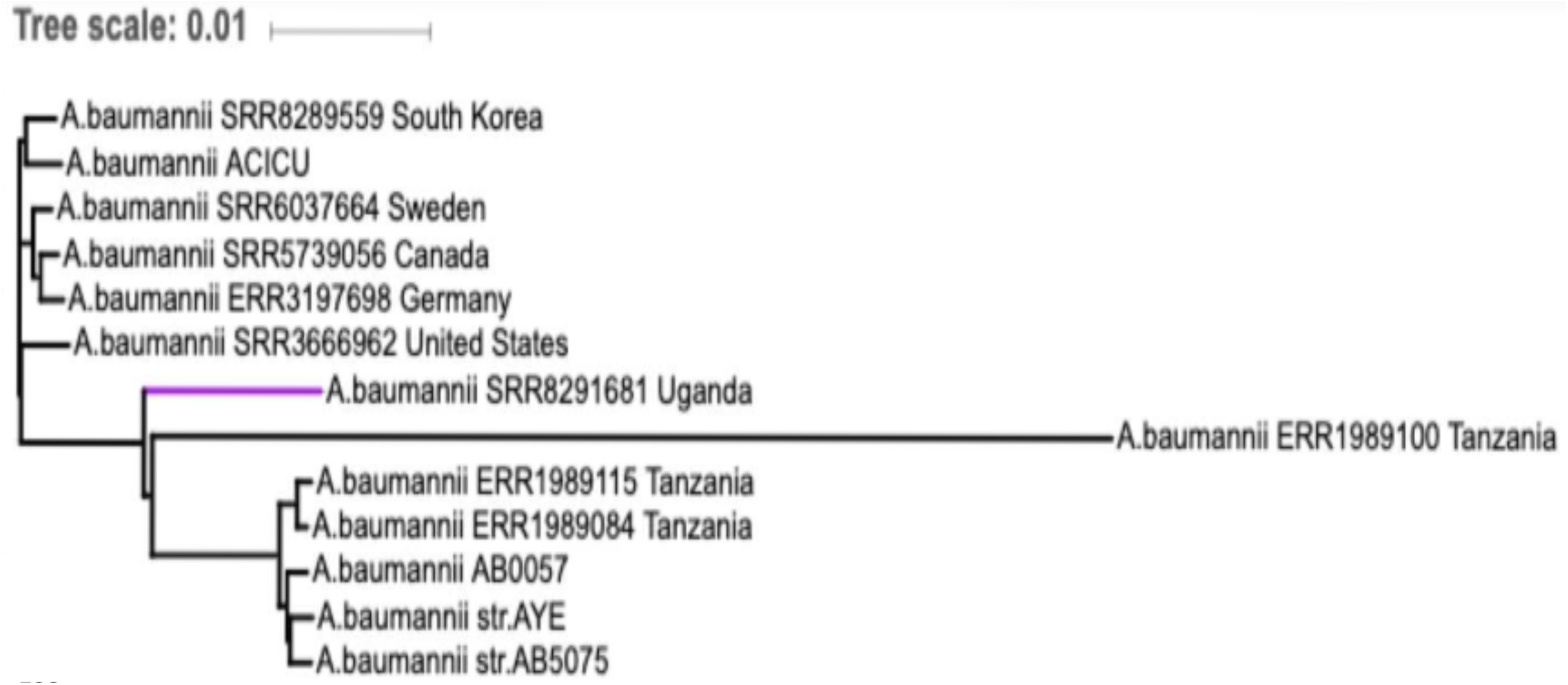
Phylogenetic tree. This shows the phylogenetic relationship between *Acinetobacter baumannii* SRR8291681_Uganda, strains from Tanzania (ERR1989115, ERR1989100, ERR1989084), those collected outside Africa (i.e. (SRR8289559_South Korea, SRR3666962_United States, SRR5739056_Canada, ERR3197698_Germany, and SRR6037664_Sweden)), as well as strains AB0057, AYE, ACICU, and AB5075.

### Genetic determinants of antibiotic resistance

To determine the genetic basis of antibiotic resistance, we interrogated the genome to identify acquired and intrinsic antibiotic resistance genes (**Table 1**). The acquired resistance genes were distributed throughout the genome (**Figure 2**). Resistance to β-lactams (i.e. penams, second-, third, and fourth-generation cephalosporins, cephamycins and carbapenems) was mediated by the previously described *Acinetobacter*-derived cephalosporinases (ADCs) and carbapenem-hydrolyzing class D β-lactamase genes (CHDLs)^5–7^. Allelic copies or mutants (i.e. *lpxA* and *lpxC)* that have previously been linked with colistin resistance were also present^8,9^. Tetracycline resistance was mediated by the previously described acquired narrow-spectrum efflux pumps, such as the major facilitator superfamily (MFS) members^10^; an allelic copy or mutant (*tetR*) that has been linked with tetracycline resistance was also identified^11,12^. Aminoglycoside resistance genes previously described were also identified^13,14^. Macrolide resistance was mediated by chromosomally encoded macrolide-specific efflux proteins previously described^15^. Resistance to trimethoprim was mediated by a previously described trimethoprim resistance gene^16,17^. Sulphonamide resistance was mediated by a previously described mobile genetic element-encoded sulphonamide resistance gene^17,18^. Phenicol resistance was mediated by phenicol resistance genes previously described^19,20^. Mobile genetic elements (i.e. plasmids) previously documented to be mobilized into a number of species of Gram-negative bacteria by co-resident conjugative plasmids were also present, these were the RSF1010 like IncQ broad-host-range plasmids^21–24^; In addition, features of previously described *Acinetobacter baumannii* plasmids (i.e. pACICU1, pACICU2, and p3ABAYE)^25,26^ that mostly occur in clinical isolates were also present, these were the chromosome (plasmid) partitioning proteins ParA and ParB^27,28^. ISFinder analysis indicated that the genome contains numerous insertion sequences (IS), the majority of the insertion sequences belonging to the IS*3*, IS*5*, IS*66* and ISLre*2* families^29^.

**Table 1:** Acquired and selected intrinsic-antibiotic resistance genes. **Antibiotic resistance in *Acinetobacter baumannii* SRR8291681_Uganda**.*Several additional intrinsic factors, such as porins and efflux pumps which may be involved in pan-drug resistance are also encoded in the genome.

| Antibiotic category | Genes associated with<br>resistance* | Genetic<br>Mechanism | Resistance<br>Phenotype |
| --- | --- | --- | --- |
| Penams, Second- and<br>Third- generation<br>Cephalosporins,<br>Cephameycins | *bla<sub>OXA-2</sub>*, *bla<sub>OXA-51</sub>*, *bla<sub>OXA-58</sub>*,<br>*bla<sub>OXA-88</sub>*, *bla<sub>OXA-317</sub>*, *bla<sub>ADC-25</sub>* | Chromosomal;<br>Mobile genetic<br>element- mediated | β-lactamase<br>producing |
| **Fourth-generation Cephalosporins** | *bla*<sub>ADC-2</sub>, *bla*<sub>ADC-25</sub>, *bla*<sub>OXA-88</sub> | Chromosomal;<br>Mobile genetic element- mediated | AmpC β-lactamase producing |
| **Carbapenems** | *bla*<sub>OXA-58</sub>, *bla*<sub>OXA-317</sub>, and *bla*<sub>OXA-51</sub> | Chromosomal;<br>Mobile genetic element-mediated | Carbapenemase-producing |
| **Aminoglycosides** | *aph*(3'')-Ib, *aph*(6)-Id, *aph*(3')-Ia, *aac*(3)-IId, *aac*(3)-IIa, *aph*(3')-I, *aph*(3'')-I, *aph*(6)-Ic, *aph*(6)-Id, *aac*(3)-II, III, IV, VI, VIII, IX, X | Mobile genetic element- mediated | Aminoglycoside resistance |
| **Macrolides** | *macA*, *macB* | Chromosomal - mediated | Macrolide resistance |
| **Tetracyclines** | *tetA*, *tetB*, *tetR* | Mobile genetic element- mediated | Tetracycline resistance |
| **Trimethoprim** | *dfrA1* | Mobile genetic<br>element- mediated | Trimethoprim<br>resistance |
| **Sulphonamides** | *sul2* | Mobile genetic<br>element- mediated | Sulphonamide<br>resistance |
| **Phenicol** | *floR* family | Mobile genetic<br>element- mediated | Phenicol<br>resistance |
| **Colistin** | *lpxA, lpxC* | Chromosomal –<br>mediated | Colistin resistance |

**Figure 2.**
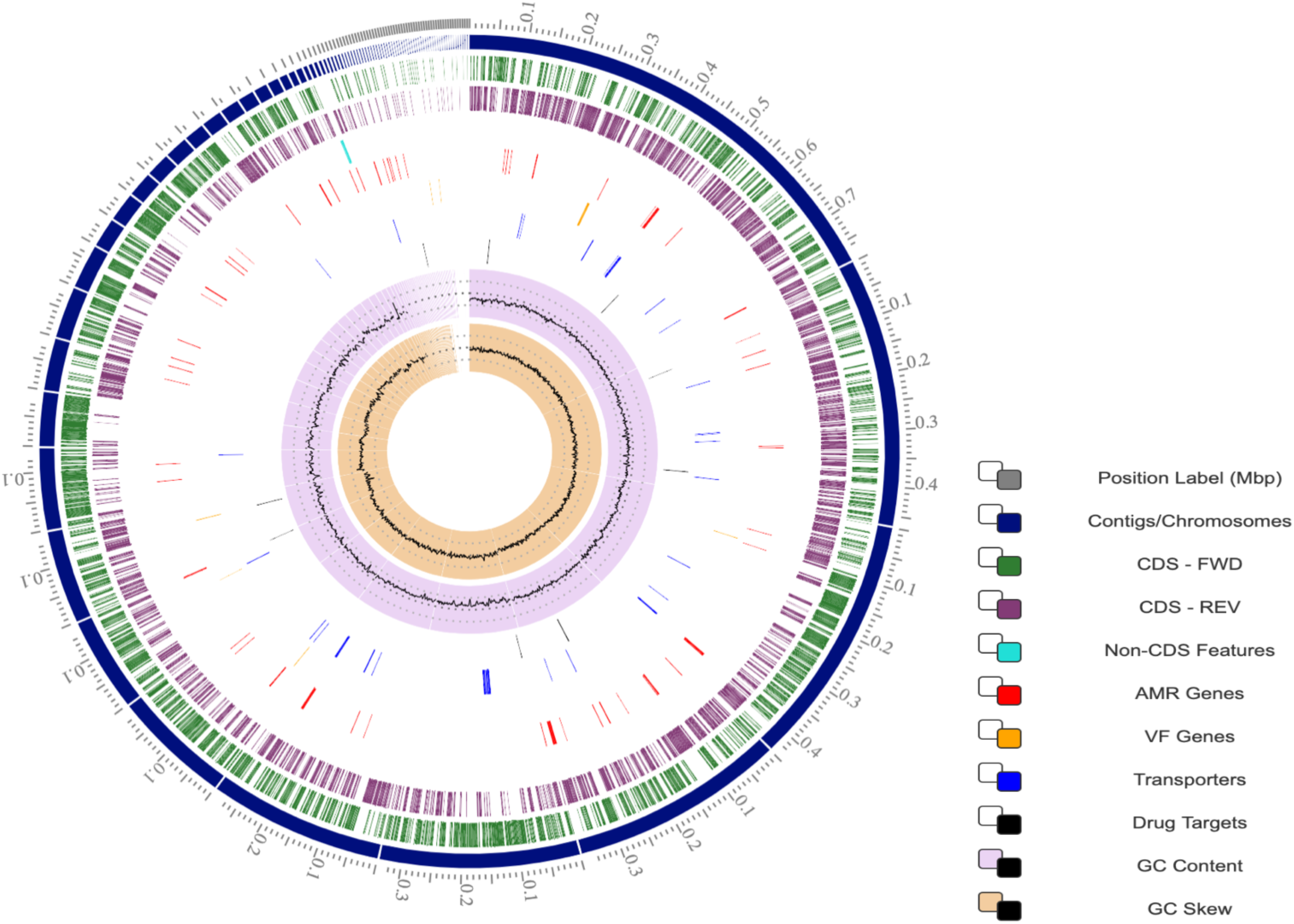
Graphical circular map of the genome of *Acinetobacter baumannii* SRR8291681_Uganda. **Circular map of the genome**. The two inner black circles indicate the G+C content plotted against the average G+C content of 38.95%; the green and purple circles show GC skew information, and the outer circles display the open reading frames (ORFs) in opposite orientations.

## Discussion

This is the first genomic analysis of a pan-drug resistant *Acinetobacter baumannii* in Uganda. The definition of this isolate as pan-drug resistant was based on the definition of the Centers for Disease Control and Prevention (CDC) and the European Centre for Disease Prevention and Control (ECDC) assessment criteria^30^

Comparable to other Gram-negative bacteria, several intrinsic and acquired mechanisms of antibiotic resistance previously described are present in *Acinetobacter baumannii* SRR8291681_Uganda^31–34^. These mechanisms can be classified into three broad categories, these include; (i) antibiotic-inactivating enzymes, (ii) reduced access to drug targets of the bacteria, attributable to the decrease in the outer membrane permeability caused by the loss or reduced expression of porins, over-expression of multi-drug efflux pumps, and (iii) mutations that change drug targets or cellular functions (i.e. alterations in the penicillin-binding proteins). These resistance mechanisms explain the evolution of antibiotic resistance of the pan-drug resistance genotype of *Acinetobacter baumannii* SRR8291681_Uganda.

Coupled with the intrinsic antibiotic resistance mechanisms, steps in the evolution of antibiotic resistance of *Acinetobacter baumannii* SRR8291681_Uganda likely followed in a sequential manner that started with the, (i) acquisition of multiple antibiotic resistance genes by the antibiotic-susceptible strain from antibiotic-resistant strains of bacteria through conjugation, transformation, or transduction in horizontal gene transfer^31–35^; genes that enabled the organism to produce enzymes with the ability to degrade antibiotics, to express antibiotic efflux pumps that prevent antibiotics from reaching their intracellular targets, to modify the antibiotics’ target sites, or to trigger the production of alternative metabolic pathways that enable the bypass of action of antibiotics^31–35^, (ii) integration of the multiple antibiotic resistance genes into the host bacteria’s genome or plasmids; and their distribution herein, facilitated by deoxyribonucleic acid transposons and bacterial insertion-sequences^36,37^, and (iii) consequent vertical gene transfer of the antibiotic resistance genes in clonal spread^38–40^.

In *Acinetobacter baumannii* SRR8291681_Uganda evolution of antibiotic resistance likely followed from penams to second- and third-generation cephalosporins along with the naturally occurring resistance to fourth-generation cephalosporins, to cephamycins, carbapenems and then to the polymyxins; along with resistance to other classes of antibiotics (i.e. aminoglycosides, tetracyclines, trimethoprim, sulphonamides, and phenicols; along with intrinsic macrolide resistance).

*Acinetobacter*-derived cephalosporinases (ADCs) precisely the extended-spectrum AmpC-type β-lactamases (ESAC) have been described to possess the ability to hydrolyze extended-spectrum cephalosporins and monobactams, this has been attributed to a Pro210Arg substitution and a duplication of an Ala residue at position 215 (inside the O-loop); however, these have also been described to have no potential of compromising the efficacy of carbapenems^41,42^. Furthermore, *Acinetobacter baumannii* has also been described to express the *bla*_*OXA-51*_, another chromosomally-encoded intrinsic β-lactamase for which many point-mutant variants have been described; these enzymes normally exhibit low-levels of carbapenemase activity and have the potential of being overproduced especially when their genes are provided with efficient promoters by insertion sequences (i.e. ISAba1 or ISAba9), making *Acinetobacter baumannii* consequently resistant to carbapenems^34,43,44^.

Over several decades, carbapenems have been reserved for the treatment of known or suspected multi-drug resistant bacterial infections including those by *Acinetobacter baumannii*^6,45^. However, over time, various acquired β-lactamases (i.e. carbapenem-hydrolyzing class D β-lactamase genes (CHDLs)) have been identified to mediate carbapenem resistance in *Acinetobacter baumannii*^6,46^. Despite the ability of these to hydrolyze carbapenems, they only show but limited ability to hydrolyze extended-spectrum cephalosporins; their association with extra intrinsic resistance mechanisms, however, has been described to provide a high level of carbapenem resistance^41^. *bla*_*OXA-58*_ and its variants have been documented to occur in worldwide isolates of *Acinetobacter baumannii*^47^. These genes have been described as plasmid-encoded, as well as being associated with specific insertion sequences which play a role in enhancing expression of the same; these have, however, been described to not be part of transposons^41,48^.

Over the years, several aminoglycoside-modifying enzymes including phosphotransferases, acetyltransferases, and adenyltransferases have been described and documented to occur in *Acinetobacter baumannii*, these mediate antibiotic resistance via reducing or abolishing binding of the aminoglycoside molecule to the ribosome^41,49^. The genes encoding these enzymes have also been documented to often occur in mobile genetic elements^41,49^.

Tetracycline resistance has been documented to occur when reversible binding of tetracycline to the 30S ribosomal subunit occurs and as a result, inhibiting protein translation^41,49,50^. In *Acinetobacter baumannii*, the *tetA* and *tetB* have been described as specific efflux pump proteins that actively remove tetracycline from the cells^41,49,50^.

The main mechanisms of resistance to polymyxins and colistin antibiotics involve the; (i) reduction of the net negative of the outer-membrane protein by modification of lipid A, an essential component of the bacterial lipopolysaccharide (LPS); and (ii) proteolytic cleavage of the antibiotic compound and exclusion of peptides by a broad spectrum efflux-pump^51^. In *Acinetobacter baumannii*, antibiotic resistance gene variants or mutants *lpxA* and *lpxC* that mediate polymyxin and colistin resistance via molecular bypass have been documented^8,9^.

Despite the significance of pan-drug resistance phenotypic data in defining true pan-drug resistance in bacterial isolates^54^, the current study was unable to obtain this data for *Acinetobacter baumannii* SRR8291681_Uganda. The study therefore described the isolate as pan-drug resistant as defined by CDC and ECDC basing on genomic analysis.

In addition, genomic analysis alone was utilized to describe mobile genetic element-mediated antibiotic resistance evolution in *Acinetobacter baumannii* SRR8291681_Uganda. This was informed by a similar study that determined the genetic basis and evolution of pan-drug resistance in bacteria by interrogating the genome to identify acquired as well as intrinsic antibiotic resistance genes^54^.

The inability to compare the differential evolution of antimicrobial resistance in this study was to the most part attributed to the fact that the only available *Acinetobacter baumannii* genomes from Africa particulary Tanzania that were added as part of the analysis had earlier been described as extended-spectrum β-lactamase encoding and had been confirmed as such post-sequencing. These genomes therefore could have been inadequate to compare with those of the carbapenem and colistin resistant *Acinetobacter baumannii* SRR8291681_Uganda.

## Conclusion

Although limited by the absence of pan-drug resistance phenotypic data neccessary in defining bacteria as truly pan-drug resistant this study provided the first genomic analysis of a pan-drug resistant *Acinetobacter baumannii* from Uganda and Africa and described mobile genetic element-mediated antibiotic resistance in the same isolate. The study provides a basis to track trends in antibiotic resistance as well as identification of emerging resistance patterns in *Acinetobacter baumannii* in Uganda. Lastly, the study recommends that continued survailance using both phenotypic and genotype data be done continuously to identify country-specific data on antibiotics effective for antibiotic resistant bacteria.

## Abbreviations

WHO: World Health Organization;
MLST: Multilocus sequence typing;
ST: Sequence typing;
ADCs: Acinetobacter-derived cephalosporinases;
CHDLs: Carbapenem-hydrolyzing class D β-lactamase genes;
MFS: Major facilitator superfamily;
IS: Insertion sequence;
CDC: Centres for Disease Control and Prevention;
ECDC: European Centre for Disease Prevention and Control;
ESAC: Extended-spectrum AmpC-type β-lactamases;
PATRIC: Pathosystems Resource Integration Center;
CARD: Comprehensive Antimicrobial Resistance Database;
NDARO: National Database of Antibiotic Resistant Organisms;
JCVI: J. Craig Venter Institute;
CC: Clonal complex;
GC: Global complex

## Acknowledgments

Not applicable.

## Author contributions

Conceptualization: DA, GM. Data Curation: DA, GM, IS. Formal analysis: DA, GM, IS. Methodology: DA, GM, IS. Validation: DA, GM, IS. Writing-Original draft preparation: DA, GM, IS. Writing-Review and Editing: DA, GM, IS. All authors read and approved the final manuscript.

## Availability of data and materials

The datasets used and/or analysed during the current study are available from the corresponding author on reasonable request. Genomic sequence data analyzed here is freely available on NCBI and is deposited under the BioProject ID PRJNA508495 (https://www.ncbi.nlm.nih.gov/sra/SRR8291681/).

## Ethical approval and consent to participate

Not applicable.

## Consent for publication

Not applicable.

